# Optimizing bacteriophage screening and isolation methods for microbial samples derived from different body sites of cattle

**DOI:** 10.1101/2025.07.04.663187

**Authors:** Gabriela Magossi, Samat Amat

## Abstract

Bacteriophages are gaining increased research attention as alternatives to antibiotics and microbiome manipulation tools to enhance feed efficiency and animal health in cattle. However, challenges associated with phage specificity, microbial ecosystem variations, and the absence of effective screening methods have hindered harnessing the power of phage application in cattle. The objectives of this study were to (i) optimize phage screening method for microbial samples obtained from different cattle body sites, (ii) isolate lytic phages against key bovine pathogens and commensal bacteria, and (iii) characterize the isolated phages and their bacterial hosts. A total of 1,214 samples from different cattle body sites (n = 1194) and environmental sources were screened using 13 phage detection methods, including one high-throughput approach. Eighty-three phages were isolated, primarily from ruminal fluid (59), feces (15), vaginal (7) and nasopharyngeal swabs (1), and fetal ruminal fluid (1). The bacterial hosts inhibited by these phages were from 29 genera, with *Bacillus* (34), *Escherichia/Shigella* (8), *Shouchella* (5), *Corynebacterium* (4), and *Lysinibacillus* (4) being the most common. No phages were identified against bovine pathogens including *Trueperella pyogenes, Mannheimia haemolytica, Pasteurella multocida, or Moraxella bovis.* Method 12 demonstrated the highest efficiency in phage recovery, particularly from ruminal samples. The isolation of phages against commensal bacteria from the gastrointestinal, reproductive, and respiratory tracts, and fetal gut highlights their potential for microbiome modulation to improve cattle health and feed efficiency. These findings underscore the need for further research into pathogen-targeting phage isolation in cattle.

**Importance:** Bacteriophage application holds a promising potential for improving feed efficiency and reducing enteric methane emissions and fighting against antimicrobial resistant bacterial pathogens in cattle. In the present study, we optimized a bacteriophage screening and isolation method that could be used to isolate bacteriophages against commensal bacteria from the ruminal, respiratory and reproductive tract microbiota in cattle. We isolated 83 phages that are lytic to bacterial species originating from rumen, vagina, nasopharynx of healthy adult cattle, as well as the intestinal tract of 6-month-old calf fetus. Overall, our study provides an important basis for the development of bacteriophage-based interventions to modulate ruminal, reproductive and respiratory tract microbiomes and thereby improve animal health and feed efficiency while reducing methane emissions in cattle.

## Introduction

Bacteriophage screening against bovine bacterial pathogens is essential due to the rising antimicrobial resistance (AMR) in pathogens associated with bovine respiratory disease (BRD) (1), liver accesses (2), mastitis (3) and bovine infectious keratoconjunctivitis (pinkeye) (4). The increasing prevalence of multidrug resistant pathogens in cattle poses significant throat to the human, animal and environmental health, highlighting the impetus need for developing alternatives to antibiotics to maintain sustainable livestock production globally. Bacteriophage therapy offers a promising alternative to antibiotics (5). Bacteriophages are viruses that specifically infect and lyse bacterial cells (6, 7). Based on life cycle, phages are categorized into lytic and lysogenic (temperate) phages, with the former being the ideal for the phage therapy due to its capacity of rapid distribution and destruction of host bacterial cells (8).

Phage therapy has been evaluated for treating or preventing infectious diseases in farm animals. For example, bacteriophages have been tested as prophylaxis and treatment for enteropathogenic *E. coli* in calves (9), piglets (10), and small ruminants (11), and showed promising efficacy in reducing mortality and morbidity in treated animals (12). The phages shed in the feces of those phage cocktail receiving animals also showed improvement of animal health after used as fecal transplantation (13). The efficacy of phage cocktails against 7 enteropathogenic *E. coli* strains in preventing and treating diarrhea in calves has also been demonstrated preciously (13). Phage therapy has also been successfully employed to control *Campylobacter jejuni* and *Campylobacter coli* (14), and to reduce mortality in chickens inoculated with pathogen *Salmonella Typhimurium* (15).

In addition to the application of phage therapy as an antimicrobial alternative to control infectious diseases in food-producing animals, recently phages have garnered increasing research interest due to their potential use in precise modulation of microbiome and host-microbiome interactions (16, 17). Phages are highly specific to their bacterial hosts and thus have minimal to no effect on the other commensal members of the microbiome where the phage host resides (18). Cattle body harbors diverse, site-specific and complex microbial communities. Microbiomes associated with gastrointestinal, reproductive and respiratory tracts, as well as oculus and mammary glands are important in maintaining overall cattle health and productivity (19, 20). Among the different microbiomes present in and on cattle body, the ruminal microbiome plays an important role in defining feed efficiency, enteric methane emissions, and overall host health and immune development (21, 22). The reproductive microbiomes (male and female) have also been increasingly known for their importance in defining reproductive health and fertility (23–26). Besides, the respiratory microbiome is also important in maintaining pulmonary health via providing colonization resistance against BRD pathogens (27–30). Thus, developing bacteriophage-based microbiome manipulation to improve feed efficiency, enteric methane mitigation, and gut, respiratory and reproductive health in cattle (31) holds a great potential in such an era where cattle production continues under pressure to improve its output while reducing the environmental impact and reliance on antibiotic usage (32).

The sampling nature, bacterial cell density, and microbiome complexity differ across the gut, reproductive and respiratory tracts in cattle. This coupled with the potential bacterial cell exchanges between different microbial ecosystems (gut, respiratory and reproductive tracts) within a cattle body highlights the necessity to optimize a bacteriophage screening method that can be used for screening different sample types with an optimal phage recovery rate. Isolating and characterizing phages from different bovine tissues, fluids, and swabs come with several challenges. These challenges could be overcome by refining the process involved with sampling, phage enrichment, and plaque assays. Classical methods of screening and isolating bacteriophages was developed early 20th Century (33)(), and the current most common method used for bacteriophage screening from environmental samples is the double-layer agar assay (34–38). The double-layer agar method has been modified, and improved over the years (33, 39). Double-layer agar-based phage screening has been applied to ruminal samples with a limited phage isolation success rate (40) despite the rumen has been believed to harbor relatively large phage population (10^9^ and 10^11^ viral particles/mL of ruminal fluid) (41) comprising more than 25 lytic morphotypes (40, 42, 43). Lytic phages targeting specific ruminal bacteria such as *Streptococcus bovis* (44–46), *Bacteroides ruminicola* (47), *Ruminococcus* (48), and *Butyrivibrio fibrosolvens* (49) have been isolated. Despite the limited number of phages having been isolated from the ruminal ecosystem, the recent metagenomic data (43) suggest significant potential for new phage discoveries within the rumen that could be used to manipulate ruminal microbiome composition, its functions and ruminal fermentation.

Thus, there is an impetus need for optimizing the current available bacteriophage methods for screening microbial samples originating from different anatomical sites of cattle (rumen, respiratory, and reproductive tracts, and fetal associated microbial samples). In the present study, we screened a relatively large number of microbial samples originating from different cattle body sites for isolation of phages that can inhibit bovine pathogenic bacteria and commensal bacterial members; We optimized phage screening method that could be used for phage screening from microbial samples obtained from different cattle body sites. In addition, we performed whole-genome sequencing and microscopic imaging on a subset of isolated phages to gain genomic and morphological insights into those bovine origin bacteriophages.

## Materials and Methods

This study involved screening of a relatively large number of samples obtained mainly from beef cattle; These beef cattle from which samples were collected were from 7 different animal trials conducted in North Dakota State University (NDSU), Fargo, ND, USA. All experimental procedures were approved by the NDSU Institutional Animal Care and Use Committee (IACUC). Animal trial #1 (Hempseed cake study): IACUC protocol ID A21010; Animal trial #2 (Lactipro study): IACUC protocol ID 20210068; Animal trial #3 (Maternal microbiome modulation and offspring calf microbiome development): IACUC protocol ID 20210043; Animal trial #4 (One carbon metabolite supplements during early gestation on offspring calf development) IACUC protocol ID A21049; Animal trial #5 (Vagino-uterine microbiome characterization between pregnant and non-pregnant cattle): IACUC protocol ID A21061; Animal trial #6 (Characterization of the ocular microbiota in pinkeye cattle): IACUC protocol ID 20220029); Animal trial #7 (Multigeneration impact of supplementation of VTM during gestation on offspring calf development and performance): IACUC protocol ID A21047.

### Brief description of the animal trials from which the samples were collected and used for phage screening

The animal samples used for bacteriophage screening in the present study originated from 7 different beef cattle trials conducted in NDSU livestock husbandry facilities and a brief description of each of these studies are presented below.

Animal trial #1 (Hempseed cake study): This study evaluated the effects of feeding 20% hempseed cake to 19-month-old angus-crossbred heifers (n = 32) compared to 20% dried distillers’ grains with solubles (DDGS) in the microbiome composition of the gastrointestinal, respiratory, and reproductive microbiome (50) These heifers were housed in NDSU Beef Cattle Research Center (BCRC).

Animal trial #2 (Probiotic study): This animal trial was conducted to evaluate the impact of supplementing a Lactipro NXT *Megasphaera elsdenii* probiotic to beef heifers fed a high-energy diet; for this trial, a total of 32 yearling heifers were used, and the animal trial was conducted at BCRC in NDSU. Ruminal fluid (n = 81) collected from the heifers in this trial were used for phage screening in the present study (Table 1).

**Table 1.** List of bovine pathogen bacterial hosts used in phage screening from animal and environmental samples.

| Pathogen species | Strain ID | Source | Species |
| --- | --- | --- | --- |
| *Trueperella pyogenes* | 306 | Liver abscess | Cattle |
| *Trueperella pyogenes* | 311 | Liver abscess | Cattle |
| *Trueperella pyogenes* | 315 | Liver abscess | Cattle |
| *Trueperella pyogenes* | 2023_3_60 | Ruminal tissue | Cattle |
| *Trueperella pyogenes* | 2022_1_14 | Ruminal tissue | Cattle |
| *Trueperella pyogenes* | 2022_1_94 | Ruminal tissue | Cattle |
| *Trueperella pyogenes* | ATCC 19411 | Lung | Swine |
| *Trueperella pyogenes* | 22_4153abc | Abscess | Otter |
| *Trueperella pyogenes* | 22_4153lg | Lung | Otter |
| *Trueperella pyogenes* | 22_2564 | Perineum | Cattle |
| *Trueperella pyogenes* | 22_2565 | Lung | Cattle |
| *Trueperella pyogenes* | 21_2041 | Lung | Deer |
| *Trueperella pyogenes* | 7762 | Lung | Cattle |
| *Trueperella pyogenes* | 22_2570 | Milk | Cattle |
| *Trueperella pyogenes* | 16_8561 | Abscess | Cattle |
| *Trueperella pyogenes* | 2645 | Abscess | Deer |
| *Mannheimia haemolytica* | 3129 | Nasopharynx | Cattle |
| *Mannheimia haemolytica* | EO_764 | Nasopharynx | Cattle |
| *Mannheimia haemolytica* | EO_700_1 | Nasopharynx | Cattle |
| *Mannheimia haemolytica* | EO_744_2 | Nasopharynx | Cattle |
| *Mannheimia haemolytica* | EO_728_2 | Nasopharynx | Cattle |
| *Mannheimia haemolytica* | EO_755_7_A | Nasopharynx | Cattle |
| *Mannheimia haemolytica* | EO_748_7_B | Nasopharynx | Cattle |
| *Mannheimia haemolytica* | EO_749_-1 | Nasopharynx | Cattle |
| *Mannheimia haemolytica* | EO_744_-1 | Nasopharynx | Cattle |
| *Mannheimia haemolytica* | EO_786_-1 | Nasopharynx | Cattle |
| *Mannheimia haemolytica* | EO_777_28_C | Nasopharynx | Cattle |
| *Mannheimia haemolytica* | EO_766_28_A | Nasopharynx | Cattle |
| *Mannheimia haemolytica* | EO_777_42_B | Nasopharynx | Cattle |
| *Mannheimia haemolytica* | EO_716_28_A | Nasopharynx | Cattle |
| *Mannheimia haemolytica* | EO_760_28_A | Nasopharynx | Cattle |
| *Mannheimia haemolytica* | EO_712_42_A | Nasopharynx | Cattle |
| *Mannheimia haemolytica* | EO_764_42_A | Nasopharynx | Cattle |
| *Mannheimia haemolytica* | EO_704_28_B | Nasopharynx | Cattle |
| *Mannheimia haemolytica* | EO_751_42_B | Nasopharynx | Cattle |
| *Mannheimia haemolytica* | EO_735_42_A | Nasopharynx | Cattle |
| *Mannheimia haemolytica* | EO_786_42_A | Nasopharynx | Cattle |
| *Pasteurella multocida* | EO_730_42_A | Nasopharynx | Cattle |
| *Pasteurella multocida* | EO_785_1_B | Nasopharynx | Cattle |
| *Pasteurella multocida* | EO_720_42_A | Nasopharynx | Cattle |
| *Pasteurella multocida* | EO_769_28_A | Nasopharynx | Cattle |
| *Pasteurella multocida* | EO_716_28_A | Nasopharynx | Cattle |
| *Pasteurella multocida* | EO_763_42_A | Nasopharynx | Cattle |
| *Pasteurella multocida* | EO_777_-1 | Nasopharynx | Cattle |
| *Pasteurella multocida* | EO_730_1 | Nasopharynx | Cattle |
| *Pasteurella multocida* | EO_739_28_A | Nasopharynx | Cattle |
| *Pasteurella multocida* | EO_716_2 | Nasopharynx | Cattle |
| *Pasteurella multocida* | EO_718_1 | Nasopharynx | Cattle |
| *Pasteurella multocida* | EO_746_28_B | Nasopharynx | Cattle |
| *Pasteurella multocida* | EO_777_28_B | Nasopharynx | Cattle |
| *Pasteurella multocida* | EO_730_42_A | Nasopharynx | Cattle |
| *Pasteurella multocida* | EO_753_28_A | Nasopharynx | Cattle |
| *Pasteurella multocida* | EO_787_2 | Nasopharynx | Cattle |
| *Pasteurella multocida* | EO_740_42_A | Nasopharynx | Cattle |
| *Pasteurella multocida* | EO_712_42_B | Nasopharynx | Cattle |
| *Pasteurella multocida* | EO_751_42_A | Nasopharynx | Cattle |
| *Pasteurella multocida* | EO_777_42_A | Nasopharynx | Cattle |
| *Pasteurella multocida* | 2873 | Nasopharynx | Cattle |
| *Escherichia coli* | BHI_cont | Environmental | Environmental |
| *Moraxella bovis* | 7116 | Ocular swab | Cattle |
| *Moraxella bovis* | 194735-2 | Ocular swab | Cattle |
| *Moraxella bovis* | 3481-2 | Ocular swab | Cattle |
| *Moraxella bovis* | 19.13095-6 | Ocular swab | Cattle |
| *Moraxella bovis* | 198338-1 | Ocular swab | Cattle |

**Table 2.**
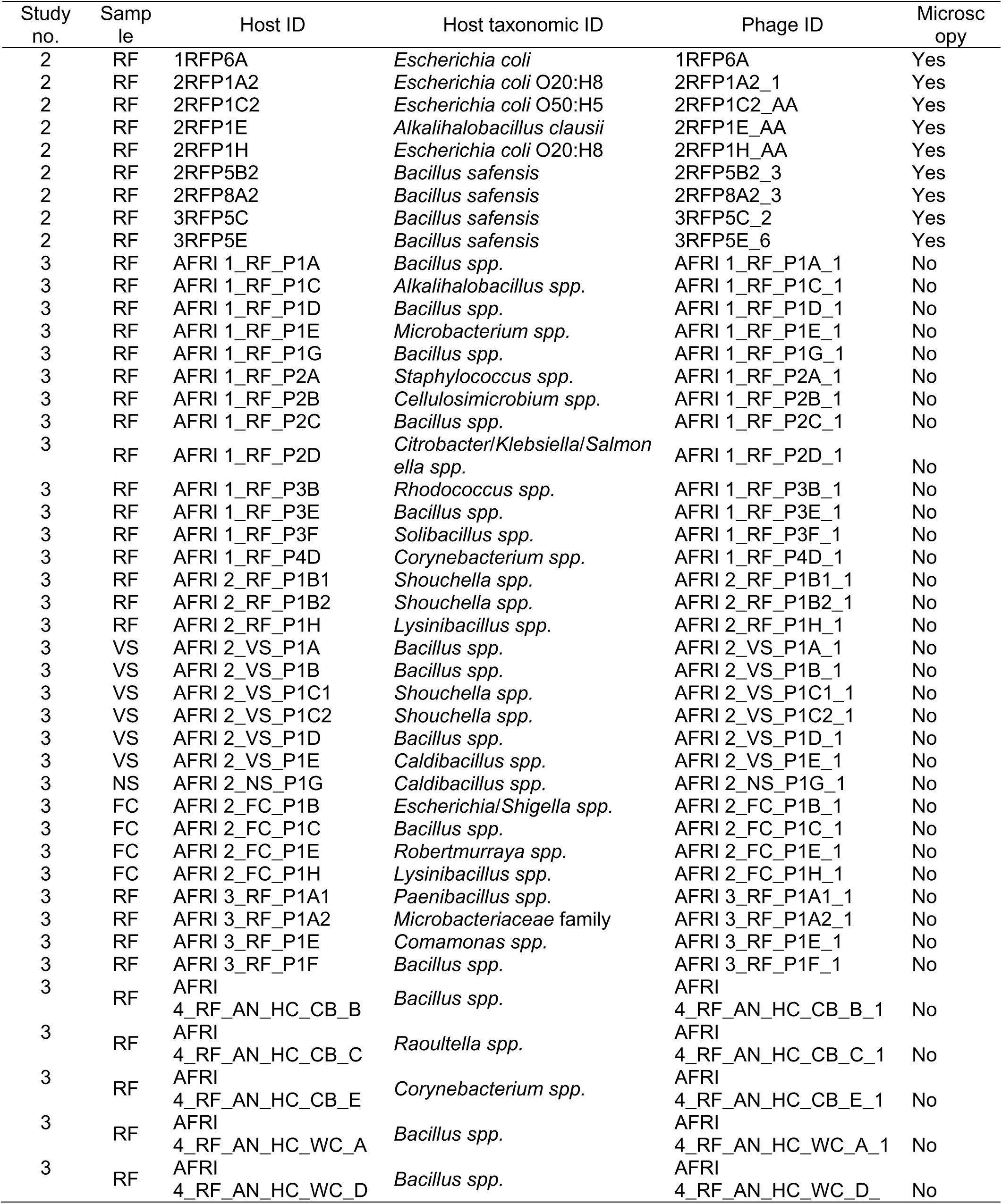
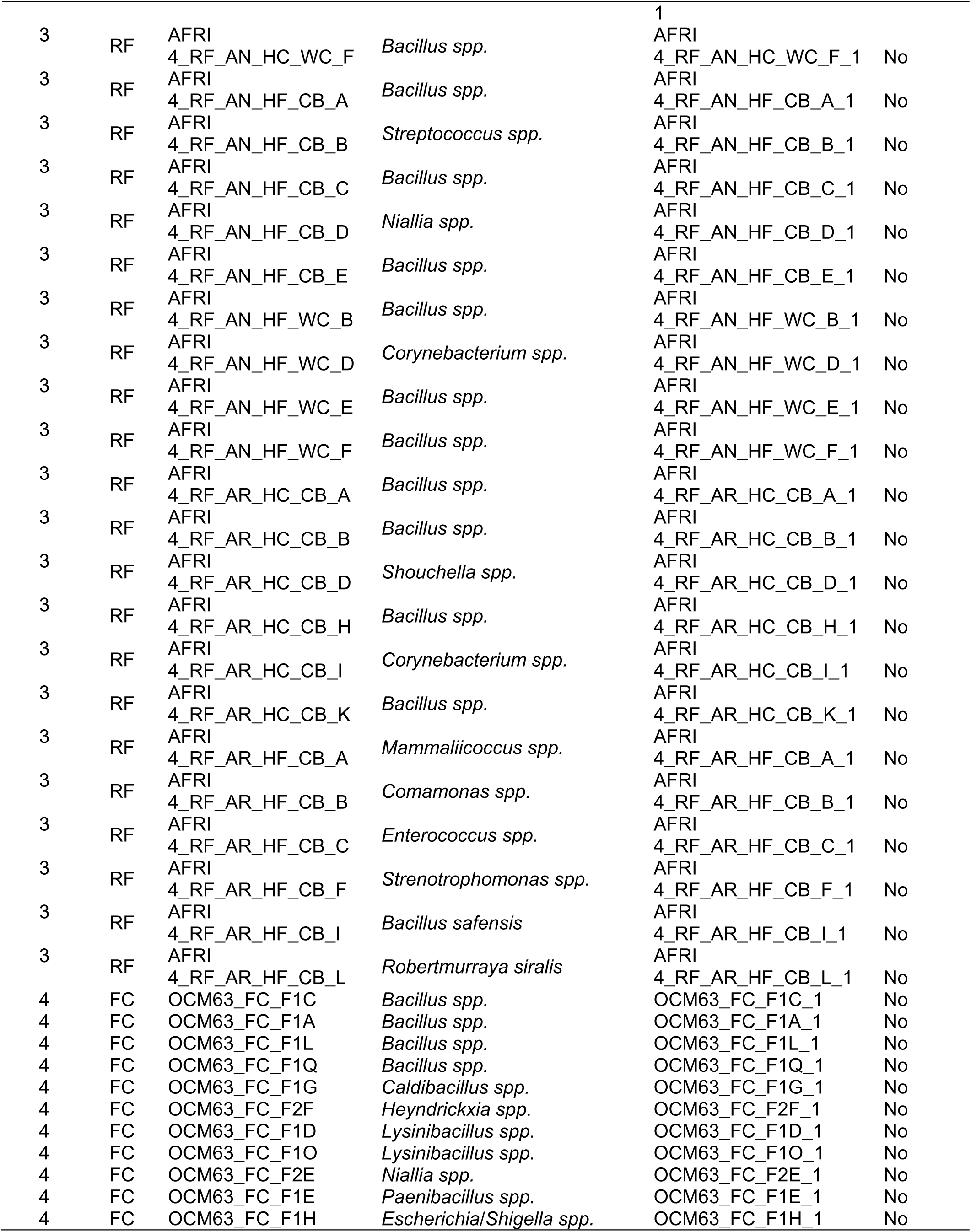
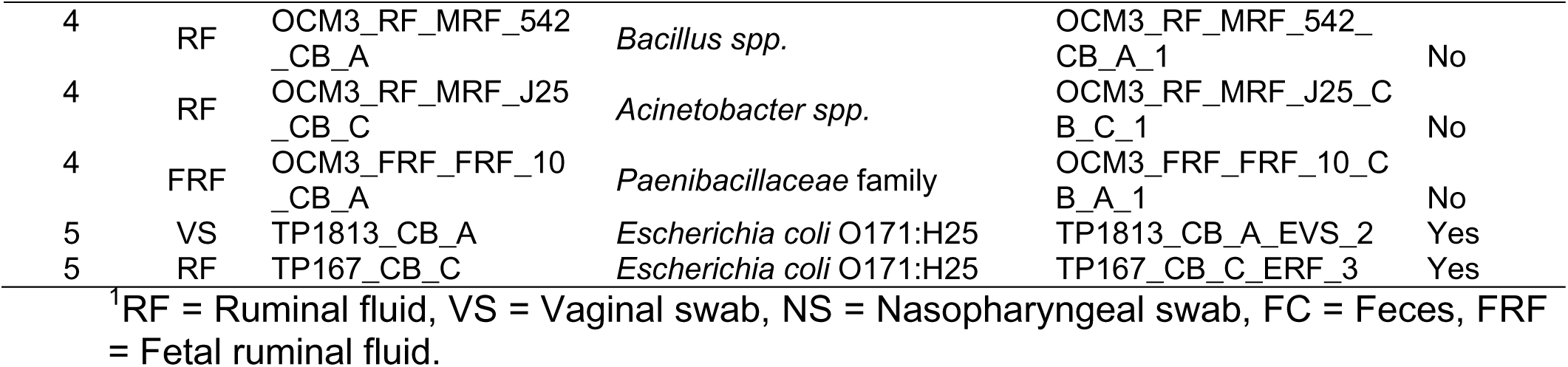
List of bacteriophages isolated from bovine samples through different studies at NDSU and their respective bacterial hosts.

Animal trial #3 (AFRIMMAT study): This animal trial was conducted to evaluate the impact of altering maternal microbiome via high forage or concentrate diets during gestation on the offspring calf microbiome development, animal performance, and enteric methane mitigation. For this trial, a total of 120 beef heifers were used. Heifers which were managed on either a high forage diet (75% forage, 25% concentrate) or a high concentrate diet (75% concentrate, 25% forage) at the NDSU BCRC (Amat et al., unpublished). Ruminal fluid (n = 89), vaginal swabs (n = 11), nasopharyngeal swabs (n = 14), and fecal (n = 4) samples were used for phage screening from this study.

Animal trial #4 (OCM study): This animal trial was conducted to evaluate the impact of one carbon metabolite (OCM) supplementation during gestation on fetal and offspring calf development. For this, a total of 32 pregnant beef heifers, which were fed either control or restricted diets, with some receiving OCM supplementation, including methionine, choline, folate, and vitamin B12 as described in our previous publication (51). This study was conducted at NDSU Animal Nutrition and Physiology Center (ANPC). The heifers were inseminated with female-sexed semen and maintained on OCM dietary treatments until 260 days of gestation, at which time they were slaughtered and tissue from dams and fetuses were collected (unpublished data). Ruminal fluid (n = 79), vaginal swabs (n = 78), nasopharyngeal swabs (n = 32), and fecal samples (n = 22) as well as 20 fetal ruminal fluid and 19 fetal meconium samples were used for phage screening from this study.

Animal trial #5 (Pregnancy efficiency study): This animal trial was involved in characterizing vaginal and uterine microbiota in cows and heifers at the time of artificial insemination to identify vagino-uterine microbiota signature associated with pregnancy (26). For this trial, Angus-crossbred cows (27 open, 31 pregnant) from which both uterine and vaginal swabs were collected at the time of artificial insemination (AI). These cows were raised in a pasture at the NDSU Central Grassland Research and Development Center (Streeter, ND, USA). Vaginal swabs were also collected from beef heifers (26 open, 33 pregnant) 2 days before AI. These heifers were housed in NDSU ANPC and as part of OCM trial (Animal trial 4). Vaginal (n = 15) and uterine (n = 15) swabs collected from this trial were used for phage screening in the present study.

Animal trial #6 (Pinkeye study): This animal trial was conducted to evaluate the ocular microbiota associated with pinkeye in cattle. Ocular swabs were collected beef cattle (n = 100) with and without pinkeye. These sampled cattle originated from NDSU Beef Cattle Research and Teaching Unit, and some cow-calf farms in North Dakota, USA. A total of 25 ocular swabs were used for phage screening in the present study.

Animal trial #7 (VTM study): This animal trial was conducted to evaluate multigeneration impact of maternal vitamin and mineral (VTM) supplementation during pregnancy on fetal and offspring calf development (52). For this trial, a total of 16 pregnant heifers were used, and they were housed in the NDSU ANPC. Ruminal fluid and vaginal swabs were used for phage screening from this study.

Of note, except for the Animal trial #6, all the cattle originated from the same cow-calf herd raised in NDSU CGREC. The cattle management at BCRC and ANPC were done with the same personnel and microbial sample collections across all these different trials were conducted by the same team.

### Sample collection for phage screening

#### Ruminal fluid (RF) sample collection

Ruminal samples were collected from angus cattle and these samples were sampled from beef steers, dams and heifers, as well as dairy cattle. The cattle were restrained in a hydraulic cattle chute and a metal speculum was placed in the mouth so that a flexible PVC tube could be passed through the esophagus and into the rumen as described in our previous publications (50, 53). The tube was worked through the ruminal mat and then a light vacuum was applied to retrieve the ruminal fluid. The ruminal fluid was collected into a clean plastic side-arm Erlenmeyer flask, gently swirled, and aliquoted into either a sterile 15- or 50-mL falcon tubes. Tubes containing ruminal fluid used for phage screening were stored in warm water, transported to the lab and processed upon arrival.

### Vaginal swab (VS) sample collection

The vaginal swab samples were collected from both dams and heifers (26, 50, 53). While animals were restrained in a mechanical chute, one or two sterile cotton tipped swabs (10 cm) were inserted simultaneously into the vagina and vaginal canal to collect as much mucous material as possible. The swab tips were broken and placed into 2 mL cryovials. Then, a combination of up to 12 swabs, originating from 6 different animals, were combined in 50 mL centrifuge falcon tubes and kept on ice and transported to the lab.

### Uterine swab (US) sample collection

The uterine swab samples were collected from angus bred cows, as described previously (26, 50, 53). Briefly, double-guarded culture swabs (Reproduction Provisions L.L.C., Walworth, WI) were inserted into the uterus, passing the cervix with the guide through rectal palpation. Swab tips were cut and placed into sterile 2 mL centrifuge tubes and kept on ice and transported to the lab.

### Deep nasopharyngeal swab (DNS) sample collection

The deep nasopharyngeal swab (DNS) samples were collected from cattle as described previously (54). Briefly, animals were restrained in a mechanical chute, DNSs were inserted into the right nostril of each animal after the outer region of the nostril was cleaned with paper towels. Swab tips were cut with 70% ethanol sterilized metal pliers, placed in sterile 1.5ml microcentrifuge tubes and stored on ice until brought back to the lab.

### Ocular swab (OS) sample collection

Ocular swab samples were collected from beef cattle infected with IBK, or healthy herd mates, across the North Dakota, on the field by instructed personnel and shipped to our lab at NDSU at different time points (55). Ocular swabs were collected from cattle diagnosed with pinkeye and healthy herd mates using Puritan Opti-Swab with Liquid Amies Collection and Transport System (Puritan, Guilford, ME, USA). For swabbing, cattle were placed in a hydraulic squeeze chute and the head was manually restrained. Before swabbing, the sampled eye was wiped with a clean paper towel sprayed with 70% paper towel to remove any debris. Then, eyelids were opened, and the conjunctival and cornea tissues were gently swabbed. Immediately after sample collection, the tips of the swabs were broken and placed in 1 ml of sterile Amies transport medium and stored on ice for transport to the lab.

### Fecal sample collection

Feces were collected manually while wearing gloves cleaned with 70% ethanol from the rectum of angus heifers and deposited into sterile whirl-pak bags. Bags were kept on ice until brought back to the lab. Then, using a sterile disposable spoon, one scoop of four different fecal samples were transferred into a 50 mL tube, next, 45 mL of Dulbecco’s phosphate buffered saline (DPBS; Corning, Corning, NY, USA) was added and the contents vortexed. Tubes were stored at 4°C until further processing.

### Liver tissue and liver abscess sample collection

Liver tissue samples were obtained via liver biopsy at the NDSU ANPC as previously described (56). Briefly, animals were restrained in a mechanical chute and clippers were used to remove the hair between the 10^th^ and 11^th^ intercostal spaces on the right side of the animal. Then, 3 mL of 2% lidocaine hydrochloride was administered as a local anesthetic and the skin cleaned by scrubbing betadine and alcohol around the incision site. Finally, a Tru-Cut Biopsy Device (Merit Medical, South Jordan, UT) was used to obtain the liver tissue sample that was immediately transferred into a sterile 2 mL cryotube, kept on ice, and transported to the lab.

The liver abscess samples were obtained from three different beef cattle, abscesses were dissected from surrounding liver tissue aseptically after the animal was slaughtered and the liver was removed from the carcass. Abscesses were then placed into sterile whirl-pak bags and shipped on ice to our lab at NDSU where they were processed.

### Fetal ruminal fluid and meconium sample collection

Fetal samples were collected from beef cattle fetuses (N = 20; 180-day-old fetuses) harvested at the time of slaughter as described previously (53). The uterus was removed and the surface cleaned with 70% ethanol, next, an incision was made with a scalpel, and the fetus removed. Fetuses’ organs were removed with their contents intact, then a 50cc syringe with an 18-gauge needle was used to extract the fetal ruminal fluid from the rumen, and a sterile scoop was used to transfer meconium into a sterile 15 mL centrifuge tube. Both samples were kept on ice and transported to the lab.

### Milk sample collection

Two milk samples were obtained from the NDSU Dairy Farm in Fargo (ND, USA). One sample was taken from the collection tank containing milk from all cows (N = 100) milked that day, and the second one was taken directly from the teat of an animal with mastitis. Milk samples were placed in sterile 120 mL plastic specimen collection cups.

### Runoff and drinking water, and wastewater sample collection

The runoff and drinking water samples were collected from the NDSU BCRC with sterile 120 mL specimen collection cups. The runoff water samples were collected from water that pooled at the lower point at the end of the animal holding pens, from the surface ground. The drinking water was collected from the drinking water tanks located in between 2 pens, from two different tanks. The wastewater samples were collected in the Fargo area. Approximately 100 mL aliquots were stored in plastic specimen cups and kept on ice and transported to the lab.

### Soil sample collection

The samples were collected using a soil sample probe tool from the soils of or around the area where beef cattle had been housed at the NDSU Beef Unit facility. The core soil was placed into sterile whirl Pak bags and kept on ice and transported to the lab and processed immediately upon arrival in the lab.

### Sample collection for the high throughput assay

Samples (n = 109) for the high throughput assay were collected from pregnant beef heifers (approx. 260 days of gestation) at time of slaughter at seven time points (sequential days, four different animals each day) from the same project looking into OCM supplementation (Animal trial#4, OCM study). These samples included DNS, OS, VS, tracheal swabs (TR), and RF. A bone saw was used to cut through the skull of each animal halfway up the nasal cavity, and the DNS were obtained by using cotton tipped swabs into the exposed nasopharyngeal cavity. The OS were acquired by using cotton tipped swabs to swab the surface of the eyes, including the inside of both the upper and lower eyelids. Additionally, cotton tipped swabs were inserted into the trachea after the organs, including the lungs, were removed from the carcasses, reaching the entirety in length of the trachea tube. Once the rumen was removed from the carcass, the organ surface was cleaned with 70% ethanol and a sterile scalpel was used to make a small incision into the rumen, then, a 50 mL tube was used to collect RF from its contents. Finally, VS were obtained as described above. The tips of the swabs were removed and placed into 2 mL cryotubes and kept on ice until transported into the lab for processing. Samples were pooled (all NS together, n = 4, all VS together, n = 4, etc.) and stored in 2 mL cryotubes containing 30% glycerol at -20°C.

### Bacteriophage host, culture conditions, and their origins

The bacterial species selected as target hosts for phage screening included pathogens of interest to cattle. These included the BRD associated bacterial pathogens *Mannheimia haemolytica*, *Pasteurella multocida,* and *Histophillus somni,* and the multifaceted opportunistic pathogen *Trueperella pyogenes*, the IBK associated pathogens *Moraxella bovis* and *Moraxella bovoculi,* as well as the potential foodborne pathogens *Escherichia coli* and *Salmonella typhimurium* (Table 1). Commensal bacteria isolated from animal samples were also used for phage screening. *T. pyogenes* isolates originated from bovine liver abscesses, *M. haemolytica*, *P. multocida*, and *H. somni* were isolated from DNS of beef cattle. *M. bovis* and *M. bovoculi* were isolated from ocular swab samples of cattle with IBK that were received by the Veterinary Diagnostic Lab (VDL) at NDSU. The *E. coli* and *S. Typhimurium* isolates were part of our culture collection and were cultured from cattle feces. All bacterial cultures were stored at -80°C in brain heart infusion (BHI; BD, Franklin Lakes, NJ) + 20% glycerol 2 mL cryotubes. They were all cultured using either tryptic soy agar (BD, Franklin Lakes, NJ, USA) with 5% defibrinated sheep’s blood (TSAb) or BHI agar and/or broth and incubated at 37°C supplemented with 5% CO_2_.

To culture and isolate commensal bacteria from the different cattle samples, 1 mL of pooled samples (RF or swabs) were serially diluted by transferring 50 µl of vortexed liquid sample into 450 µL of DPBS (Dulbecco’s phosphate-buffered saline; Thermo Fisher Scientific, Branchburg, NJ, USA). Next, 100 µL of dilutions 10^-1^ and 10^-2^, for swabs, and 10^-4^ and 10^-5^, for RF, were spread plated on TSAb plates and incubated at 37°C with 5% CO_2_ overnight. After incubation, the colonies with visibly different morphologies within the plate, and between plates were picked and sub-streaked onto new Columbia blood (CB; BD, Franklin Lakes, NJ, USA) agar plates for isolation using a sterile disposable needle. Plates were then incubated at 37°C + 5% CO_2_ overnight. The overnight grown isolates were transferred into 1 mL of BHI + 20% glycerol in 2 mL cryotubes and stored at -80°C.

### Methods for phage detection from cattle and environmental samples

#### Sample filtering

A cheesecloth was used to separate larger particles from the RF, then, RF from individual animals were pooled together (4-8 animals per pool), mixed, transferred into 50 mL centrifuge tubes, and centrifuged at 3,900 *x g* for 15 min at 4°C. Next, the supernatant of this first centrifugation was transferred to 2 mL microcentrifuge tubes, and centrifuged at 20,000 *x g* for 15 min at 4°C. Finally, the supernatant was filtered using 0.22 µm syringe filters, and the filtrate was used in the downstream processing.

### Swab sample processing

Vaginal swab tips stored in microcentrifuge or cryotubes were pooled together (from 4 to 8 swabs per pool) in a sterile 15 mL centrifuge tube containing 10 ml of sterile DPBS and thoroughly vortexed (for minimum 20 s). One mL of swab suspension pools was used for culturing while the remaining supernatant was filtered using 0.22 µm syringe filters and 10 ml syringes, then, 100 µL of the filtrates were transferred into 5 mL of BHI broth containing 100-200 µL of a target bacterial isolate culture or pool of up to 5 isolates cultures and incubated at 37°C shaking at 220 rpm overnight. This was considered as an enrichment step. After incubation, the enrichment tubes were centrifuged at 15,000 x *g* for 15 min at 4°C and the supernatants filtered using 0.22 µm syringe filter.

### Polyethylene glycol (PEG) precipitation

To readily concentrate phage particles that might be present in large volumes of liquid media, a 20% PEG6000 (w/v) + 2.5M sodium chloride (NaCl) solution was added to the liquid at a 5:1 ratio (v/v) of sample filtrate to PEG + NaCl solution (57, 58). Then, the mixture was wrapped in aluminum foil and incubated at 4°C for 48 h. After 48 h, the solution was transferred into 2 mL microcentrifuge tubes, and centrifuged at 12,000 x *g* for 30 min at 4°C. The supernatant was discarded, and the pellet was resuspended using 100 µL of SM buffer (50mM Tris-HCl [pH7.5], 100mM NaCl, 8mM MgSO_4_, 0.01% gelatin; G-Biosciences, St. Louis, MO), and stored at 4°C until further use.

### Preparing soft agar

The hard agar, or regular solid agar, was prepared by adding 1.5% (w/v) of agar to the broth media and pouring onto sterile plastic petri dishes. The hard agar plates were used for normal bacterial culturing and used as the bottom agar plates for the soft agar overlay essays.

Soft agar was prepared by adding 0.5% (w/v) of agar to BHI broth prior to autoclaving, then, 3 mL of soft agar was poured in sterile test tubes. In both agar preparations, hard and soft, the media was cooled to 55°C in a water bath after being taken out of the autoclave, then, Calcium (CaCl_2_) and Magnesium (MgSO_4_) salts were added to a final concentration of 2.5mM.

### Spot assay on double-layer agar

For the double-layer agar spot assay, 100 µL of overnight bacterial host culture were added to 3 mL of BHI soft agar, mixed, then poured on top of a regular solid BHI agar plate. Plates were swirled to cover the entire surface with the soft agar top layer and let to solidify on the benchtop at room temperature. Once the soft agar top layer was solidified, 10 µL of concentrated filtrates were spotted on the surface using a P20 pipette and tips with enough space between spots on the plate. Spots were then left to dry before plates were inverted and incubated at 37°C with or without 5% CO_2_ overnight. After incubation of spot assay plates, areas of bacterial lysis that were identified by a clearing in bacterial lawn, either as a complete circular clearing or as plaques, were scraped using a 10 µL loop and placed into 1 mL sterile DPBS, vortexed vigorously, then centrifuged at 12,000 x *g* for 5 min to pellet cell debris and agar. Supernatants were then filtered using a 0.22 µm syringe filter into new 1.5 mL microcentrifuge tubes. To determine the best dilution for double agar, a spot dilution assay was used. For spot dilutions, the filtrates obtained from the spot assay were serially diluted by 1:10 using sterile DBPS until the 10^-10^ dilution, then, 10 µL of each dilution was spotted onto a newly made double-layer agar plate, inoculated with 100 µL of overnight bacterial host culture. Plates were left to dry on the benchtop at room temperature before they were inverted and incubated at 37°C with 5% CO_2_ overnight.

### Phage stock preparation

The double-layer agar assay was done by adding 100 µL of spot filtrate or phage stock to a 1.5 mL tube containing 100 µL of bacterial host overnight culture. This mixture was left to incubate at room temperature for 15-20 min for phage adsorption, then transferred into 3 mL of BHI soft agar, mixed, and poured on top of a regular BHI agar plate, swirling the plate as to cover the entire surface with the soft agar top layer. Plates were let to solidify at room temperature before being inverted and incubated at 37°C with 5% CO_2_ overnight. After incubation, plates with between 30-300 plaques were selected for calculating phage stock titters in PFU/mL. Isolated plaques were transferred into 500 µL of SM buffer, filtered using a 0.22 µm syringe filter and stored at 4°C.

### Phage enrichment and serial enrichments

Sample enrichments and serial enrichments were accomplished by inoculating broth with target bacterial host and adding 1 mL or 1 g of samples to the mixture. For a single enrichment, 50-200 µL of overnight bacterial culture in BHI broth were added to 5 ml BHI broth in 15 ml centrifuge tubes and incubated at 37°C shaking 100 rpm overnight. 50 µL of each was added when more than one isolate was used to inoculate into the broth. When a single isolate was used, then 100 µL of culture was added. For those bacteria with longer latent growth phases, 200 µL of culture were added to the broth. After incubation, tubes were centrifuged at 3,900 x *g* for 15 min at 4°C then supernatants were filtered using a 0.22 µm syringe filter and stored at 4°C. For serial enrichment, 1 mL of the unfiltered supernatant from the first enrichment step was added to a new 15 ml centrifuge tube containing 5 mL of BHI broth and the bacterial inoculum and incubated at 37°C shaking at 100 rpm overnight. This was repeated four times, after the final incubation, then the tubes were centrifuged at 3,900 x *g* for 15 min at 4°C and supernatants were filtered using a 0.22 µm syringe filter.

### Bacterial inoculation

To prepare bacterial inoculums, a new culture was started from cryopreserved glycerol stocks in 5 mL of BHI broth and incubated at 37°C with the shaking speed at 220 rpm overnight. One hundred µL of overnight culture was used to inoculate enrichment tubes and 3 mL soft agar containing tubes.

### Bacteriophage screening

The initial phage screening assays used were meant to target specific bovine pathogens described previously as bacterial hosts (Table 1), but no phage was detected against any of the pathogenic strains screened through any of the screening methods used. Therefore, we decided to start culturing commensal bacterial from the samples themselves and using those isolates as the host targets for phage screening using the same samples where they were isolated from. For example, processed RF filtrates were used as the source samples to screen for phages against the bacteria isolated from the same RF pools. The techniques and protocols used to screen phages using animal, water, or soil samples were adapted to improve on challenges and unsuccessful results obtained, culminating in a final protocol that was successful in discovering phages from animal samples when commensal bacteria strains were used as the host targets for the phage screening. All the methods and the gradual improvement procedures are described below. Of note, to improve clarity and facilitate the comprehension of the procedures and flow of the description of each step, we decided to use “Method”. The different methods listed below may reflect the changes involved in the addition of different bacterial hosts, or addition or removal of a step involved in enrichment or centrifugation, or the addition of different sample types to be screened. The different method described below does not always refer to the significant changes made in the fundamental principles or in major steps of a phage screening method.

### Optimization of bacteriophage screening methods

#### A summary of the initial different methods applied (Methods 1–11)

A series of initial methods (Methods 1–8) were tested to isolate bacteriophages targeting bovine pathogens, particularly *Trueperella pyogenes*. In Method 1, ruminal fluid (RF) samples collected from hempseed-fed cattle trial were processed via centrifugation and filtration and then screened using a double-layer agar spot assay against *T. pyogenes*. Spots were applied using both a 96-needle replicator and pipette at varying volumes (5–50 µL), but no phage activity was observed. The filtration of RF, however, proved particularly challenging due to its high viscosity and digesta content which often resulted in clogging or rupturing filters. To overcome these issues, Method 2 employed a concentrating pipette system to process larger volumes of RF filtrate. Despite this adjustment, filtration inefficiencies remained, and no lysis was detected. In Method 3, both wastewater and RF samples were used, with soft agar overlays supplemented with CaCl₂ and MgSO₄ to enhance phage adsorption. However, this also failed to produce plagues. To improve phage detection sensitivity, we introduced an enrichment step in which samples were incubated overnight in BHI broth with host bacteria prior to plating in Method 4. Both RF and wastewater samples were tested but no phages were detected from these samples. In Method 5, we screened liver biopsy samples obtained from healthy beef cattle against *T. pyogenes*. Despite T*. pyogenes* being one of the primary liver abscess-associated pathogens, the liver tissue samples didn’t yield any phages. In Method 6, we expanded the screening to DNS samples and added bovine respiratory disease (BRD) pathogens, *Mannheimia haemolytica* and *Pasteurella multocida*, to the host panel. Again, despite overnight enrichments, no phage-induced lysis was detected. In Method 7, we incorporated polyethylene glycol (PEG) precipitation to concentrate viral particles from RF prior to enrichment and plating. Although this reduced sample volume, it did not yield detectable phage activity. Method 8 explored ultracentrifugation to simplify RF processing, but did not improve clarity or throughput, and was therefore abandoned.

A substantial methodological shift was made in Method 9, where we transitioned from targeting known pathogens to using commensal bacteria isolated from the same sample pools as candidate phage hosts. Using ruminal fluid samples from heifers fed with a *Megasphaera elsdenii* probiotic supplement, we cultured and isolated a diverse set of bacterial strains and subsequently used them in screening assays. This approach led to the first successful phage detection. The bacterial host was identified as *Escherichia coli* and the phage as a member of the Caudoviricetes class. Building on this success, we introduced an enrichment step prior to plating in Method 10, using the same commensal isolates and RF filtrates from Method 9. Thismethod resulted in isolation of 8 phages against commensal bacterial host strains. After successful isolation of some phages against commensal bacterial host strains, In Method 11, we made additional change to the enrichment step, and instead of including a single host isolate, we included inoculum of a cocktail of bacterial isolates in enrichment step. Between 3-6 different isolates from the same bacterial species were used in the same enrichment tube to increase the number of possible hosts while using the same amount of sample. The bacterial target hosts were bovine pathogens *M. haemolytica*, *P. multocida*, *T. pyogenes*, *M. bovis*, and *M. bovoculi*. The source of cattle samples used were expanded from just RF to include DNS, VS, US, OS, and liver abscess material pooled together in 5-6 samples of the same type per pool, filtered and precipitated with the PEG6000 + NaCl solution. Although this method saved time and resources as compared to testing each sample for an individual bacterial host isolates, it did not result in detection of any phage against the pathogenic bacterial hosts tested.

### Method 12

For this method, serial enrichments were used instead of a single enrichment step, as used in previous methods. This change was made to enhance the phage detection rate as the initial sample volume was relatively small and thus there might be low phage cell numbers in the sample screened. The same samples used in the previous method (Method 11) (RF, DNS, VS, US, OS, and liver abscess) were screened again in this method plus the addition of soil samples.

In addition to the pathogen hosts (*M. haemolytica*, *P. multocida*, *T. pyogenes*, *M. bovis*, and *M. bovoculi*), commensal isolates obtained from the different samples were also used as target hosts for the serial enrichments. For the known bacterial hosts (pathogen hosts), their growth phase was determined by measuring the optical density (OD_600_) over time during incubation at 37°C. Then, pathogen inoculums were used when their cell growth was at the logarithmic phase for the serial enrichments and double-layer agar assays. The original samples were combined in pools as described previously prior to culturing, and these pools were added to the first enrichment tube without any filtering being done prior to it, and 250 µL of chloroform was added to the 5 mL BHI enrichment broth (only for the first enrichment).

The serial enrichment method proved to be the optimal method among all methods that we tested for the discovery and isolation of phages from animal and soil samples against commensal bacterial isolates. Chloroform was added during the first step to reduce the bacterial load of the sample being used, since a filtration step was not present, however, this addition could potentially be detrimental to the phages present in the sample, therefore this chloroform addition step was removed. All other steps were kept as before. Using this updated procedures in Method 12, bovine fetal samples were tested for the first time. The fetal samples included fetal ruminal fluid and meconium. Commensal bacteria were cultured from these samples, and the serial enrichment protocol used to screen for phages.

### High-throughput phage screening assay

In addition to the 12 different methods described above, a high-throughput screening protocol enabling the phage screening of multiple samples at the same time was also tested. This method was adapted from Olsen et al. (2020) (59). For this, different samples including DNS, OS, VS, TR, and RF were screened. Upon arrival at the lab, NS and TR swabs were pooled and transferred to a tube for each sample type containing 20 mL of DPBS, while VS and OS samples were pooled each in a tube containing 10 mL of DPBS. All tubes were vortexed thoroughly, centrifuged at 3,900 x g for 15 min at 4°C, then supernatants were transferred into 2 mL centrifuge tubes, centrifuged at 20,000 x g for 10 min at 4°C. The supernatants were filtered using 0.22 µm syringe filters, concentrated using PEG6000 + NaCl solution, and stored in 2 mL cryovials containing 500 µL glycerol + 1000 µL sample filtrate concentrate.

The selected bacterial hosts used as the targets of the high throughput phage screening were *T. pyogenes*, *M. haemolytica*, *P. multocida*, and *M. bovis*. Additionally, positive control was used to determine the efficacy of the protocol by spiking a sample with a known *E. coli* phage and testing that sample against an *E. coli* isolate. Culture negative samples were also used as negative control.

Samples were added first in each well (a different sample per well) of a 2 mL 96-well plate, the initial volume of samples (frozen concentrate filtrates) used varied from 500-1000 µL. Then, 37 µL of CaCl_2_ (0.5mM), 37 µL of MgSO_4_ (0.5mM), 636 µL of BHI broth 4.4X, and 90 µL of bacterial host culture were added to each well, except for positive and negative controls. After adding all components, BHI 4.4X was added to complete the volume of wells with 500 and 750 µL of samples, matching the level of the ones with 1000 µL sample volume, then the contents of each well were mixed by pipetting up and down before the plate was sealed with a foil film. Plates were incubated at 37°C + 5% CO_2_ overnight. After incubation, the contents of each well were transferred into 2 mL microcentrifuge tubes using a multichannel pipette, then centrifuged at 15,000 x g for 2 min at 4°C. The supernatants were filtered with a 0.22 µm syringe filters into new 2 mL deep well 96-well plate, sealed with film, and stored at 4°C until used.

A total of 200 µL of overnight host bacteria broth culture were mixed into 6 ml of soft BHI agar and poured on top of a rectangular petri dish (127.76 mm x 85.48 mm) containing TSAb hard agar and another containing BHI hard agar, covering the entire surface, and left to solidify before the next step. Filtrates were then spotted using a 96-needle applicator on both BHI and TSAb plates by immersing the applicator in the 96-well plate and gently pressing it on the soft agar overlay. The applicator was sterilized with 100% ethanol and flamed between each use and plates were incubated at 37°C overnight.

### Phage and bacterial host genome sequencing

#### Bacterial host DNA extraction

The genomic DNA from selected bacterial host isolates was extracted as previously described (60) for whole genome sequencing using a DNeasy blood and tissue kit (Qiagen, Inc., Hilden, Germany) from broth cultures. Additionally, genomic DNA was extracted from bacterial phage hosts using a Quick-DNA Fungal/Bacterial Miniprep Kit (Zymo Research, Irvine, CA, USA) following the manufacturer’s instructions, with minor modifications (61), for taxonomic identification.

### Phage DNA extraction

Phage DNA was extracted using the Norgen Biotek phage DNA extraction kit (Norgen Biotek, Thorold, ON, CA) following the manufacturer’s instructions with a few modifications as previously described (62). A pre-treatment step was performed to remove bacterial genetic material from the samples prior to extracting phage DNA. Briefly, 450 µL of filtered supernatant containing phage was treated with 50 µL of DNase I (1 U/µL; Qiagen) and 1 µL of RNase A (10 mg/mL; Qiagen) and incubated for 90 min at 37°C. Next, 20 µL of Ethylenediaminetetraacetic acid (EDTA) at a concentration of 20 mM was added with the goal of inactivating the activity of the DNase and RNase enzymes, and lastly, 1.25ul of Proteinase K (20 mg/mL) was added and the mix was incubated at 56°C for 90 minutes. For the extraction of genomic DNA from the 1RFP6A and 3RFP5C_6 phages, however, a bead beating lysis method was employed prior to genomic DNA extraction with the Zymo Quick-DNA Miniprep kit (Zymo Research Corporation, Irvine, CA, USA).

### DNA concentration and quality assessment

Next, a Nanodrop spectrophotometer (Thermo Fisher Scientific, Waltham, MA) was used to assess the quality of the extracted genomic DNA samples, and a Qubit 4 fluorometer (Thermo Fisher Scientific) with the double-stranded DNA (dsDNA) high-sensitivity assay kit (Thermo Fisher Scientific) was used to quantify the extracted genomic DNA.

### Sanger sequencing and taxonomic identification of bacterial hosts

Sanger sequencing of the 16S rRNA gene was used for the taxonomic identification of bacterial host isolates. A PCR reaction containing 10 µL of iQ Supermix (Bio-Rad Laboratories Inc., Hercules, CA), 1 µL of 27F primer (5′-AGAGTTTGATCMTGGCTCAG -3′; 10 µM), 1 µL of 1492R primer (5′-TACGGYTACCTTGTTACGACTT -3′; 10 µM), 2 µL of DNA template, and 6 µL of nuclease free water (Corning, Corning, NY, USA) to complete a reaction volume of 20 µL was used to amplify the near-full-length 16S rRNA gene. The cycling conditions were as follows: an initial denaturation step of 95°C for 5 min; followed by 35 cycles of 95°C for 45 s, 50°C for 30 s, and 72°C for 2 min; and a final elongation step at 72°C for 5 min. This was done in an Eppendorf Mastercycler (Eppendorf, Hamburg, Germany). After amplification, PCR products were sequenced by the Molecular Cloning Laboratories (MCLAB, San Francisco, CA), and sequences were identified with the Basic Local Alignment Search Tool (BLAST) and the non-redundant National Center for Biotechnology Information (NCBI) nucleotide database.

### Whole-genome sequencing and annotation

#### Phages

Libraries were preprepared for all phage genomic DNA using the Qiagen FX DNA library preparation kit (Qiagen, Germantown, MD, USA), except for phages 1RFP6A and 3RFP5C_6 which had DNA libraries prepared with the Illumina DNA Prep kit incorporating UDI 10 bp indices (Illumina, Inc, San Diego, CA, USA). An Illumina MiSeq instrument was used for phage genome sequencing utilizing the MiSeq v2 reagent kit (500 cycles; Illumina), as for the phages 1RFP6A and 3RFP5C_6, libraries were loaded into an Illumina NovaSeq 6000 instrument equipped with an S4 flow cell (300 cycles).

For all phages, sequences quality control and adapter trimming were accomplished with the Trim Galore v. 0.6.10 (63) (https://github.com/FelixKrueger/TrimGalore) tool, and genomes were assembled using Shovill v. 1.1.0 (https://github.com/tseemann/shovill) pipeline. Then, the quality of the assemblies was evaluated using QUAST (64) and annotation conducted through Pharokka v. 1.1.0 (65). Additionally, genomes were screened for the presence of virulence factors using the bacterial virulence factor database (VFDB) (66), antimicrobial resistance genes through the comprehensive antibiotic resistance database (CARD) (67), and for the presence of tRNA genes with the assistance of the tRNAscan-SE on-line resource (http://trna.ucsc.edu/tRNAscan-SE/) (Lowe & Chan, 2016). Additionally, the formula used to determine the sequencing coverage was coverage = (number of reads × read length)/genome length.

Each phage had its genome compared to those available in the NCBI database using the BLASTn tool for taxonomical identification. The combination of percent identity and percent coverage were used to define the species, genus, family, or class of the phages based on their sequence similarity (% identity x % coverage). Higher sequence similarities resulted in better classification for the phages, considering sequence identities of higher or equal to 95% to a species level, between 70%–95% to the genus level, or less than 70% to the family level (68).

### Bacterial hosts

Genomic DNA libraries from bacterial hosts were prepared using an FX DNA library preparation kit (Qiagen) and sequenced in an Illumina MiSeq instrument (Illumina) with the MiSeq v.2 reagent kit (500 cycles; Illumina) following the manufacturer’s instructions. The fastp v.0.23.2 (69) tool was used to do quality filtering and adapter trimming of sequence reads, additionally, reads that had a mean Phred quality score smaller than 15 with a sliding window of 4 bp and/or were shorter than 100 bp were removed from further analysis. Next, genomes were *de novo* assembled with SPAdes v.3.15.5 (70) with the isolate option activated. The BBMap v.38.96 aligner (https://sourceforge.net/projects/bbmap) (71) was used to remove contigs that were shorter than 500 bp using the minlength parameter configured to 500 bp. Next, contigs completeness and possible contamination were assessed with using the lineage-specific workflow of CheckM v.1.2.0 (72). Finally, QUAST v.5.2.0 (64) was used to assess assembly qualities and the Prokaryotic Genome Annotation Pipeline (PGAP) v.2022-10-03.build6384 (73) from NCBI was used to annotate the genomes. The GTDB-Tk v.2.1.1 classify workflow with the Genome Taxonomy Database (GTDB) release 207 was used to define each bacterial genome taxonomically (74, 75). All parameters for software or bioinformatics tools were set to default unless otherwise specified.

### Transmission electron microscopy imaging of phage cells

A subset of isolated phages (n = 11) was subjected to TEM imaging. Phage cell suspensions were prepared by propagating phages on BHI double-layer agar plates containing a lawn of bacterial cell host. A 10 µL loop was used to streak spread a phage stock solution onto the soft agar overlay prior to incubation at 37°C + 5% CO2 overnight. After incubations, the areas where bacterial lysis were observed on the overlay soft agar were removed and added to 500-1000 µL of nuclease free water (Corning, Corning, NY, USA), centrifuged at 12,000 *x g* for 20 min at 4°C, the supernatants were then filtered with 0.22 µm syringe filters, and filtrates submitted for imaging at the NDSU Electron Microscopy Center (Fargo, ND, USA). A drop of sample was placed on a 300-mesh formvar-carbon coated copper grid (Ted Pella Inc., Redding, California, USA) for two minutes and wicked off with filter paper. The negative stain phosphotungstic acid 0.1% was added to the samples, the pH was adjusted to 7-8, then stained samples were dropped onto the grid and allowed to stand for two minutes and then wicked off and the grid allowed to air-dry. High-resolution analytical TEM was accomplished using a JEOL JEM-1400Flash transmission electron microscope (JEOL USA, Peabody, Massachusetts, USA) at a direct magnification of 50,000 *x g*, HV = 120kV, and a NANOSPTRT15 camera with 800 (ms) exposure x 2 standard frames, gamma of 1, no sharpening, and normal contrast.

## Results

### Overview of the samples screened for phage and their origin

A total of 1,214 samples collected from 7 different beef cattle trials (98.4%), and the environment were screened for the presence of phages against bovine pathogenic bacteria and commensal bacteria originated from cattle gut, respiratory and reproductive tracts, and bovine fetuses in the present study (Table 3). Those cattle-origin samples were consisted of ruminal fluid (n = 799), feces (n = 30), and deep nasopharyngeal (n = 130), vaginal (n = 124), uterine (n = 15) and ocular swabs (n = 34), and liver biopsy tissue from healthy beef cattle (n = 10), liver abscesses (n = 3) from beef cattle, and raw milk from dairy cattle (pooled; n = 2), and soil cores (n = 12), runoff water from beef cattle facility (n = 2), beef cattle drinking water (n = 2), and municipal wastewater (n = 4).

**Table 3.** Distribution of bovine and environmental samples used for phage screening, the number of phages isolated, and the discovery rate per sample type.

| Sample type | Total number of samples | Bacteriophages | % discovery |
| --- | --- | --- | --- |
| Ruminal fluid | 799 | 59 | 7.3% |
| Feces | 30 | 15 | 50.0% |
| Vaginal swab | 124 | 7 | 5.6% |
| Fetal ruminal fluid | 20 | 1 | 5.0% |
| Nasopharyngeal swab | 130 | 1 | 0.8% |
| Drinking water | 2 | 0 | 0% |
| Fetal meconium | 19 | 0 | 0% |
| Liver abscess | 3 | 0 | 0% |
| Liver tissue | 10 | 0 | 0% |
| Milk | 2 | 0 | 0% |
| Ocular swab | 34 | 0 | 0% |
| Runoff water | 2 | 0 | 0% |
| Soil | 9 | 0 | 0% |
| Soil + feces<sup>1</sup> | 3 | 0 | 0% |
| Tracheal swab | 8 | 0 | 0% |
| Uterine swab | 15 | 0 | 0% |
| Wastewater | 4 | 0 | 0% |

<sup>1</sup>Soil + feces samples originated from the soil mixed with bovine feces inside the pens where animals were housed.

### Evolution and assessment of the phage screening methods

The use of commensal bacterial isolates as target hosts and the inclusion of an enrichment step to the phage screening process were considered as the first and second successful modifications from the initial tests, respectively. Therefore, a further improvement on the enrichment step was made for Method 12, and this improvement was performing four rounds of serial enrichment. In this method, both bovine bacterial pathogens and commensal bacterial strains were used as phage hosts, and a total of 67 phages were isolated. All of these phages inhibited the commensal bacterial hosts tested. No phage against pathogenic bacterial hosts was detected.

From the 2 runoff water, 2 drinking water, 30 fecal, 28 DNS, 62 RF, 12 soil, and 1 VS samples were screened by this method, 16 phages from feces, 1 from DSN, 45 from RF, and 5 from VS samples were isolated (Table 3).

Serial enrichments of sample filtrates were found to be the most effective procedure that enhanced the phage detection rate from both cattle and environmental samples. However, the addition of chloroform in the first enrichment round of the four serial enrichments used was later recognized as unnecessary and that it may compromise the phage detection outcomes. The original purpose of adding chloroform in this early step of enrichment was to lysate bacterial cells to release possible phage particles from the infected host cells (intracellular), and to reduce the unrelated bacterial cell load present in the screened samples. However, chloroform could also degrade and reduce the stability of some phages, in addition to increasing the quantity of cell debris from lysed bacterial cells. Therefore, we removed this step involved in chloroform addition during the enrichment step. By this modified method (Method 12), 2 more phages from the RF (n = 20 screened), and one phage from the fetal RF (n = 20) were recovered. The 19 fetal meconium samples screened did not yield any phage detection. No phages were detected against any bovine pathogens tested in this method.

### High-throughput screening method

For the high-throughput 96-well plate method, there was a total of 109 samples used, including 22 RF, NS, TR, and VS, and 21 OS samples, collected in seven different days. Sample filtrates were used in soft agar lawns inoculated with the *M. haemolytica*, *P. multocida*, *M. bovis*. No lysis was observed against any of the tested pathogens.

Overall, among the different phage screening methods that we used, Method 12 was found to be the optimal assay that enabled us to identify phages from different sample types obtained from differ cattle body sites against ruminal and other commensal bacteria. After removing the chloroform addition step during enrichment, the updated Method 12 was chosen as the most effective method for screening different microbial samples obtained from different cattle body sites including ruminal fluid, fecal, vaginal and nasopharyngeal, and fetal intestine for phage isolation against commensal bacteria. This method involves 4 enrichment steps.

### Isolated phages and their respective bacterial hosts

From the 1, 214 samples screened for phages, through all of the different tested methods (13 different methods) described in this study, a total of 83 phages were isolated and banked, the equivalent of 6.8% of total samples screened. Moreover, 59 RF samples (7.4% phage recovery rate from samples screened), 15 of 30 fecal samples (50%), 7 VS (5.6%), 1 DNS (0.8%), and 1 fetal RF (5%) resulted in phage recovery (Table 3). The percentage numbers are only an overall view and any inference done over it should be interpreted cautiously, because there were different methods used, different bacterial hosts (pathogens vs. commensals), and the number of samples pooled together for phage screening. However, fecal samples followed by RF samples showed the highest phage recovery rate (Table 3). Overall, most of the phages isolated in the present study were lytic against bacteria species within the *Bacillus* genus, making up 41% of total phages isolated (Table 4). The second most common bacterial host was *Escherichia/Shigella* (9.6%), followed by *Shouchella* spp. (6%), *Corynebacterium* spp., and *Lysinibacllus* spp. (4.8%), and *Caldibacillus* spp. (3.6%) (Table 4). However, we were not able to identify any phages against bovine pathogens such as *T. pyogenes*, *M. haemolytica*, *P. multocida*, and *M. bovis*.

**Table 4.** Bacterial host taxonomic classification and number of phages isolated per taxa.

| Host bacterial taxa | No.<br>phages | Per.<br>Phages |
| --- | --- | --- |
| *Bacillus spp.* | 34 | 41.0% |
| *Escherichia/Shigella spp.* | 8 | 9.6% |
| *Shouchella spp.* | 5 | 6.0% |
| *Corynebacterium spp.* | 4 | 4.8% |
| *Lysinibacillus spp.* | 4 | 4.8% |
| *Caldibacillus spp.* | 3 | 3.6% |
| *Comamonas spp.* | 2 | 2.4% |
| *Niallia spp.* | 2 | 2.4% |
| *Paenibacillus spp.* | 2 | 2.4% |
| *Alkalihalobacillus spp.* | 2 | 2.4% |
| *Microbacteriaceae family* | 2 | 2.4% |
| *Robertmurraya spp.* | 2 | 2.4% |
| *Acinetobacter spp.* | 1 | 1.2% |
| *Cellulosimicrobium spp.* | 1 | 1.2% |
| *Citrobacter/Klebsiella/Salmonella spp.* | 1 | 1.2% |
| *Enterococcus spp.* | 1 | 1.2% |
| *Heyndrickxia spp.* | 1 | 1.2% |
| *Mammaliicoccus spp.* | 1 | 1.2% |
| *Paenibacillaceae family* | 1 | 1.2% |
| *Raoultella spp.* | 1 | 1.2% |
| *Rhodococcus spp.* | 1 | 1.2% |
| *Solibacillus spp.* | 1 | 1.2% |
| *Staphylococcus spp.* | 1 | 1.2% |
| *Strenotrophomonas spp.* | 1 | 1.2% |
| *Streptococcus spp.* | 1 | 1.2% |
| Total | 83 | 100% |

### Taxonomic characterization of bacterial hosts and whole-genome sequencing of phages and their respective bacterial hosts

A total of 11 phage genomes were characterized by whole-genome sequencing. These phages were isolated from bovine rumen (n = 10) and vagina (n = 1). They were lytic phages, and their bacterial hosts were *Alkalihalobacillus clausii* (n = 1), *Bacillus* safensis (n = 4), and *E. coli* (n = 6). The genome results for these 11 phages have been reported as Genome Announcement (76). Briefly, the genome sizes of these bacteriophages showed considerable variation, ranging from 20,216 base pairs (bp) to 168,609 base pairs (bp) and the GC content (%) varied from 33.94-44.35. Analyzing the genomes, it was found that none contained CRISPR (Clustered Regularly Interspaced Short Palindromic Repeats) sequences, tmRNAs (transfer-messenger RNAs), virulence factors, or antimicrobial resistance genes (76). The absence of *tmRNAs*, antimicrobial resistance genes, and virulence factors ensure there are no risks of transferring these pathogenic traits to host bacteria or other members of the microbiome once phages are replicated. All the phages sequenced were tailed phages and belonged to seven different taxonomy classifications: *Caudoviricetes* (n = 3), *Felixounavirus* spp. (n = 2), *Tequatrovirus* spp. (n = 2), *Vequintavirus* spp. (n = 1), *Salasmaviridae* family phage (n = 1), *Herelleviridae* family phage (n = 1), and *Siophivirus* spp. (n = 1).

### Morphological characterization

Phages were characterized by TEM and phage cells displayed diverse morphologies, but all imaged phages were within the *Caudoviricetes* class of tailed phages and had icosahedral heads (Fig. 2). The phages displayed contractile tails as shown in Fig. 2.I and Fig. 2.J, where the phages TP1813_CBA_EVS_2 and TP167_CB_C_ERF_3 can be seen with their elongated and contracted tail. Through TEM, it is also possible to observe the similar size and structures of phage TP1813_CBA_EVS_2, isolated from the vaginal swab, and phage TP167_CB_C_ERF_3, obtained from the ruminal fluid of heifers. The phage particle sizes ranged from ∼180 to 260 nm, with head sizes ranging from ∼40 to 100 nm and tail lengths were ∼100 to 210 nm. The high contrast in TEM allowed for the visualization of fine structural details, such as tail fibers and sheath striations, providing insight into the diversity and morphology of these bacteriophages.

**Figure 1.**
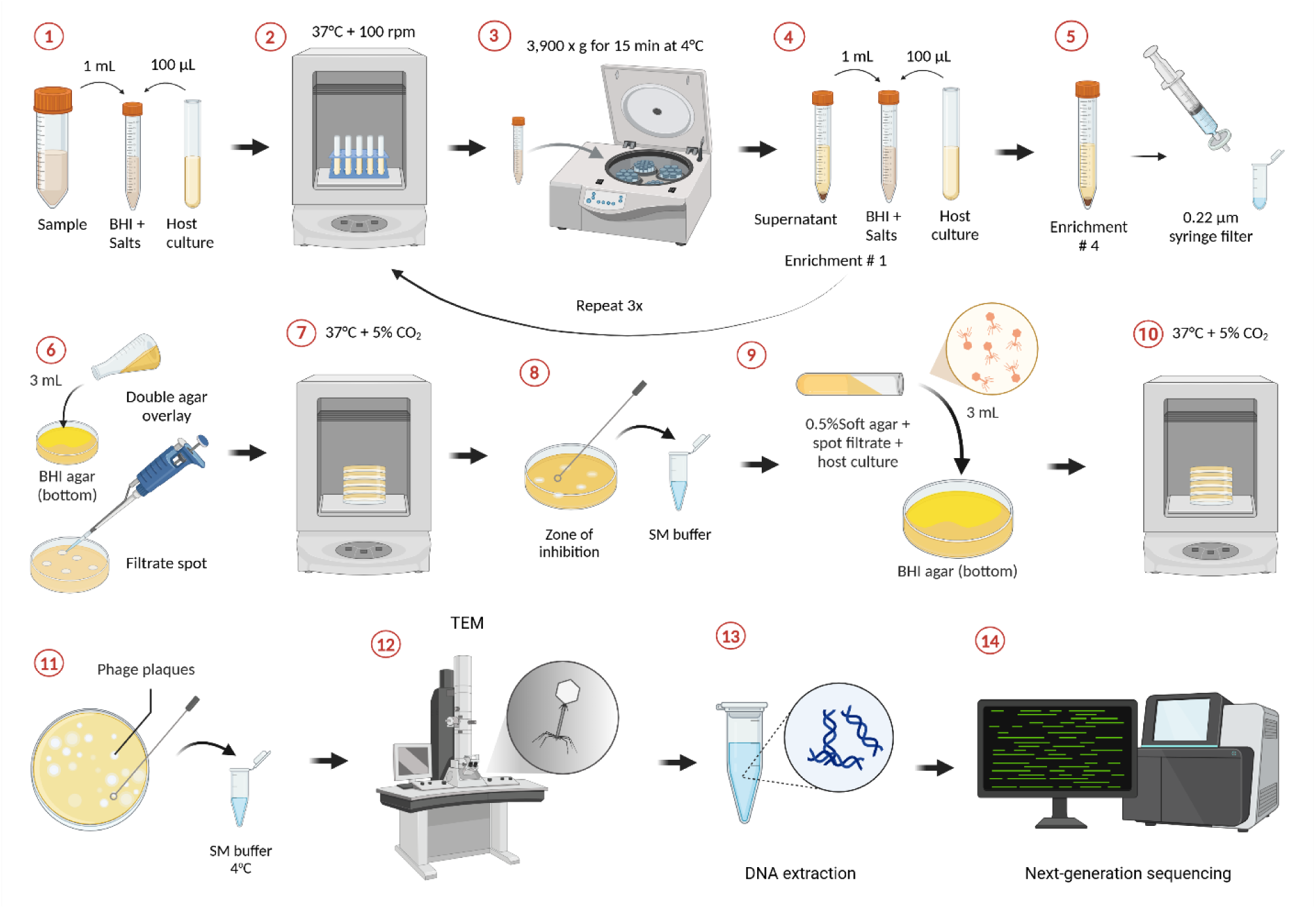
Optimized phage screening method (Method 12) using serial enrichments from animal origin samples. 1) Initial enrichment using initial sample and target host bacteria in BHI broth with the addition of CaCl_2_ and MgSO_4_. 2) Incubation step in a shaking incubator set to 37°C and 100 rpm. 3) Centrifugation step following the incubation step where centrifuge tubes are spun at 3,900 x g for 15 min at 4°C. 4) Centrifuge tubes removed from the centrifuge are used for following enrichment where supernatants and target host bacteria are combined in BHI broth with the addition of CaCl_2_ and MgSO_4._ Once centrifugation is done, the first enrichment is complete, and steps 2-4 are repeated 3 more times for a total of 4 enrichments. 5) After the fourth enrichment is complete, supernatants are filtered with a 0.22 µm syringe filter into 2 mL tubes kept at 4°C. 6) Double-layer agar spot assay. 7) Incubation of spot assay plates at 37°C + 5% CO_2_ (CO_2_ was optional). 8) Zone of inhibition due to lytic activity of phages in the filtrates picked with a loop and dispensed into SM buffer for a presumptive phage stock. 9) Double-layer agar assay to obtain isolated plaques from a mixture of 0.5% soft agar + bacterial host culture (100 µL) + presumptive phage stock from the spot assay (100 µL). 10) Incubation of double agar assay plates at 37°C + 5% CO_2_ (CO_2_ optional). 11) Selection and transfer of isolated plaques into SM buffer for long term storage at 4°C. 12) Transmission electron microscopy (TEM) of phage particles. 13) Genomic DNA extraction from isolated phages. 14) Whole genome sequencing and analysis of phages. Created in BioRender. Magossi, G. (2025) https://BioRender.com/l91n181

**Figure 2.**
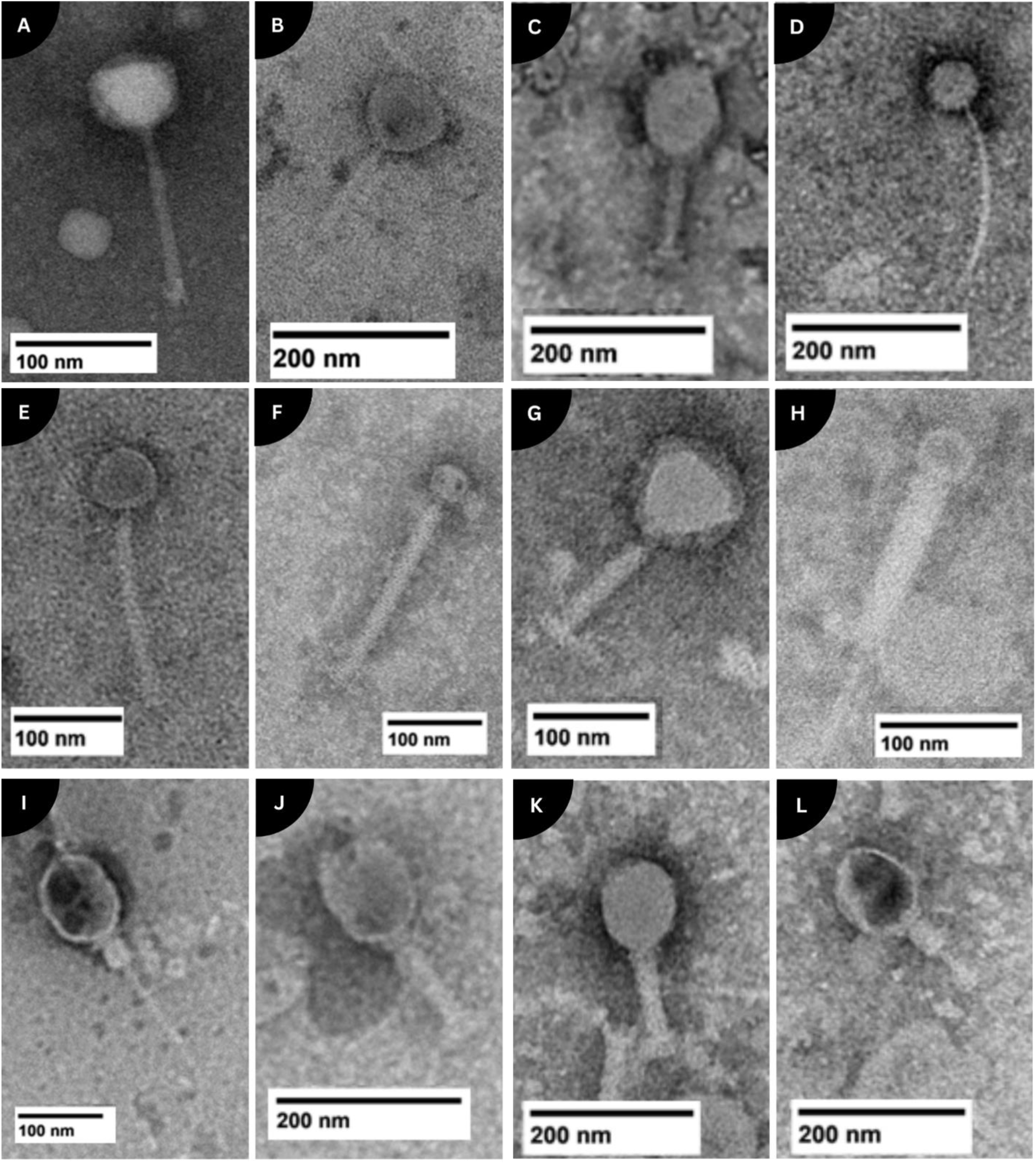
Transmission electron microscopy images of phages (host). A) 1RFP6A (*E. coli*), B)2RFP1H_AA (*E. coli)*, C) 2RFP1A2_1 (*E. coli)*, D) 2RFP1E_AA (*A. clausii*), E) 2RFP5B2_3 (*B. safensis*), F) 2RFP8A2_3 (*B. safensis)*, G) 2RFP1C2_AA (*E. coli*), H) 3RFP5C_2 (*B. safensis)*, I -J) TP1813_CB_A_EVS_2 (*E. coli*), and K-L) TP167_CB_C_ERF_3 (*E. coli)* isolated from animal origin samples with scales adjusted to nm.

## Discussion

In the present study, we screened a total of 1,194 samples obtained from different cattle body sites, including the gut, respiratory and reproductive tracts, and fetal gut, as well as environmental sources (n=20), using 13 phage detection methods. Through incremental modifications, such as the addition or removal of steps in the protocol, incorporating bacterial host samples, or expanding the variety of sample types to be screened, we optimized a phage screening method that allowed us to isolate 83 phages against commensal bacteria. Overall, with Method 12, the highest number of phages was recovered, and thus displayed the potential to be the most suitable method for screening phages from bovine gut, respiratory, reproductive, and fetal-associated microbial samples.

Among the different bovine and environmental samples screened, a greater number of phages were recovered from the ruminal fluid samples screened (n = 75). The rumen is a complex ecosystem harboring bacteria, methanogenic archaea, fungi, viruses and protozoa (41). Viral cells including bacteriophages are relatively high in quantity in the rumens (41). The ruminal phages can interact with bacteria cells in the rumen either through lysogenic or lytic cycles, and their population in the rumen can be influenced by ruminal bacterial compositions (77) and fermentation characteristics (VFA, pH), as well as diet (78). Despite the rumen being reported to contain a relatively high bacteriophage cell population (41), the phage isolation rate per sample screened was lower than that of fecal samples. This might be due to several factors. One of which is associated with the ruminal bacterial hosts used in the present study. These bacterial hosts were isolated under aerobic culturing conditions given the challenge associated with conducting phage screening procedures under a strict anaerobic condition. The anaerobic nature of the rumen may have limited in identifying and isolating even furthermore phages from the ruminal fluid samples screened.

In the present study, we isolated phages that are lytic to commensal bacterial species within *Bacillus*, *Escherichia/Shigella*, *Shouchella*, *Corynebacterium*, and *Lysinibacillus* genera originated from the cattle rumen. *Bacillus* species, such as *Bacillus licheniformis* and *Bacillus subtilis*, have been explored as probiotic supplements in cattle(79, 80)- as these *Bacillus* spp. can form spores and be able to survive in the anaerobic environment of the rumen (81, 82). Additionally, some *Bacillus* spp. have been shown to inhibit the growth of certain pathogens (83–85), and thereby improve the gut health. Furthermore, *Bacillus* species have been reported to improve feed efficiency by enhancing starch and fiber digestibility (86–90). We isolated phages against ruminal *Bacillus* spp. in the present study, and the impact of modulation of the ruminal *B*acillus population using the bacteriophages we isolated on ruminal microbiome composition, fermentation and feed digestion deserves further research.

*Shouchella* spp. and *Escherichia/Shigella* spp. primarily contribute to protein metabolism, can hydrolyze cellulose and produce H_2_, but have not been directly linked to methanogenesis (91, 92), although their interactions within the ruminal microbiome may influence overall fermentation dynamics. Some of the *Escherichia/Shigella*, and *Citrobacter/Klebsiella/Salmonella* species have been reported to be negatively correlated with ruminal fermentation (93, 94). Thus, the phages that we isolated against *Escherichia/Shigella* are particularly relevant, as these bacterial genera contain pathogenic strains and have been negatively correlated with ruminal fermentation efficiency. Reducing their populations in cattle rumen through phage administration could mitigate their negative influence on ruminal fermentation and feed digestion. Members of the *Shouchella* genus, namely *Shouchella clausii*, formerly known as *Bacillus clausii* (95), have been used as probiotics in humans (96), similarly to *Enterococcus* spp. and *Streptococcus* spp. (97). Although *Corynebacterium* genus does encompass pathogenic species such as mastitis-associated *C. amycolatum* and *C. pseudotuberculosis* (98), certain *Corynebacterium* species are associated with amino acid fermentation and nutrient degradation in the rumen (99). *Lysinibacillus* is one of the a least characterized genera in the ruminal microbiota *Lysinibacillus fusiformis* has been linked with increased the feed digestibility and ruminal VFA production and reduced methanogens by limiting H_2_ availability to methanogenic archaea (100). Given that these bacterial species are positively associated with increased nutrient utilization and reduced methanogenesis, the phages that we isolated against these commensal bacterial species should be investigated thoroughly. The purpose of ruminal phage application is to induce changes in ruminal bacterial populations and thereby enhance ruminal microbiome-mediated nutrient utilization and mitigate ruminal methane production. Future studies should focus on isolating pro-methanogenic ruminal commensals, such as *Ruminococcus*, which are H₂ producers that can supply methanogens with substrate, in addition to archaea viruses targeting methanogenic archaea. The targeted inhibition of these pro-methanogenic bacteria using phages could provide a novel strategy for reducing methane emissions from cattle while maintaining optimal ruminal fermentation. The optimized Method 12 in the study holds a promising method for isolating phages against both pro-methanogenic bacterial species and those negatively associated with feed efficiency in cattle.

We successfully isolated phages from vaginal and nasopharyngeal swab samples, though the isolation rate was lower compared to fecal and ruminal fluid samples. This is not commonly reported in the literature, except for a few findings, where respiratory phages have been isolated from the human throat (101) and lysogenic phages targeting vaginal bacteria could be induced, as well as reports of phage therapy against reproductive infections in cattle, have been shown (102, 103), however, direct isolation of lytic phages from the nasopharyngeal, vaginal, or uterine microbiomes of cattle remains largely unexplored. Similarly, the presence and role of bacteriophages in the bovine respiratory microbiome remain under-characterized.

Given the important role of the nasopharyngeal microbiome in maintaining respiratory health and airway immune response, particularly in high-risk cattle for BRD, microbiome modulation using bacteriophage presents a potential strategy for reducing BDR disease susceptibility (104, 105).

Likewise, the female reproductive tract microbiome significantly influences cattle health and fertility (25, 106, 107). Specific bacterial groups within the vaginal and uterine microbiomes have been associated with reproductive success or dysfunction, suggesting that microbiome-targeted interventions could enhance reproductive efficiency (26, 106). Understanding the role of phages in shaping these microbial communities within cattle urogenital tract could open new avenues for improving reproductive health in cattle (108).

A limitation encountered in this study that could have compromised the phage discovery from DNS and VS samples was that samples were collected via swab. The swab-based sample collection has its limitation as the sample volume would be small compared to ruminal fluid or feces. This would result in a reduced chance of phage detection as these swab samples may have a very limited number of phage particles. Despite this limitation, we were still able to recover phages from the nasopharynx and vaginal canal of cattle. Future studies should modify the sample collection method to ensure a significant biomass is obtained from the vagina and nasopharynx. One alternative method for nasopharyngeal sampling could be a suction pump involved sampling. A bronchoalveolar lavage sampling is also worth testing for phage screening against BRD pathogens. As for the reproductive tract, a vaginal brush-based sampling could enhance phage detection rate. We believe that the four round of serial enrichment steps we applied in Method 12 increased the chances of phage discovery from DNS and VS samples. Thus, increasing the number of enrichment steps even further for the low microbial biomass samples such as DNS and VS should be considered by future studies.

We successfully isolated a phage from the ruminal fluid of a 180-day old bovine fetus, making, to the best of our knowledge, the first report of phage isolation from healthy bovine fetuses. This discovery provides further evidence challenging the long-held sterile womb theory (105), stating that the gravid uterus and fetus are not colonized yet by microorganisms has been challenged recently, with recent evidence suggesting that the calf intestine may be colonized by bacteria and archaea prenatally (19, 53, 109). Our isolation of a bacteriophage from the calf fetal intestine suggests the presence of prenatal viral colonization in calf intestine. This is a novel observation that requires further verification by future studies.

Despite screening a relatively large number of samples (n = 1,214) and successfully isolating phages against commensal bacteria using Method 12, we were unable to identify phages targeting key bovine phages (*T. pyogenes*, *M. haemolytica*, *P. multocida*, and *M. bovis*). Several ecological and technical factors could have attributed to the unsuccessful phage isolation against pathogenic bacterial hosts.

One of the reasons could be associated with the origin of the bacterial hosts used in this study. These pathogenic strains originated from different animals involved in animal trials or from samples submitted to the veterinary diagnostic lab. These pathogenic strains have undergone multiple freeze-thaw cycles and had been cultured in the lab several time prior to being used in the phage screening. All of which might have an influence on their phage susceptibility. Thus, to increase the success rate of isolating phages against these bovine pathogen species, fresh isolation of the pathogenic isolates from the same herd of cattle, or ideally to isolate pathogenic strains from the same type of samples to be screened for the phages should be considered by future studies. The strains used in the present study may not be optimal for phage screening, and their genomes may contain anti-phage defense systems and defense islands (110). Thus, it is necessary to perform genotyping and whole genome sequencing analysis of these bovine pathogenic species before using them as bacterial hosts for phage screening. In addition, comprehensive growth curve of the pathogenic bacterial strains should be performed to identify the growth stage that is most susceptible stage for phage infection (111).

No phages to date have been identified that can infect the liver abscess and mastitis-associated pathogen *T. pyogenes* or the pinkeye-associated pathogens *M. bovis* and *M. bovoculi*. Prophages, often referred to as temperate phages, have been discovered within the genomes of *M. haemolytica* (112, 113) and *P. multocida* (114), which have the potential to be triggered into a lytic cycle, classified as *Siphoviridae* and *Myoviridae*. Temperate phages can significantly contribute to bacterial pathogenesis, virulence, and pathogen evolution through horizontal gene transfer (115). *M. haemolytica* and *P. multocida* are opportunistic pathogens that are present in the the upper respiratory tract of healthy cattle A limited number of lytic phages have been found for *P. multocida*, including a type-specific one for capsular type D; however, this phage is highly specific, only lysing non-toxigenic type D *P. multocida* and failing to infect any other strains of *P. multocida* (116). Another phage targeting *P. multocida* exhibited broader activity against capsular types B:2 and A:1(117, 118). This illustrates the specificity of type-specific *Pasteurella* phages and underscores the challenges associated with sourcing phages for this pathogen from environmental samples. For the future phage screening aimed at BRD bacterial pathogens, a thorough assessment of various types of each *M. haemolytica* and *P. multocida* should be employed to enhance the phage identification. This also indicates that using phage therapy for BRD pathogens may be difficult, necessitating a cocktail of phages for each potential type. In addition to these biological factors, limitations associated with swab-based sampling could also contribute to the unsuccessful detection of phages against screened respiratory pathogens. The opportunistic pathogen *T. pyogenes*, used as host for the phage screening of VS and an inhabitant of the reproductive tract of cattle, may have been in low abundance in animals sampled in this study because the animals were healthy without active infections. This further reduced the chances of detecting the pathogen and any associated phages. In summary, the interplay of low pathogen abundance in healthy animals, the limitations inherent in the sampling techniques employed, and the specific health status of the cattle sampled all contribute to the lack of identified phages targeting the pathogens present in the upper respiratory and reproductive tracts.

Screening bacteriophages from environmental or clinical samples presents numerous challenges, particularly when isolating unknown phages. Unlike working with stock phage solutions, which bypass many initial hurdles, successful plaque formation depends on multiple variables. Sampling limitations, along with factors related to screening methodologies, may have an impact on the success of phage screening. A few of these factors include the concentrations of Ca and Mg ions and of soft agar (36), choice of gelling agent in the soft agar overlay (119), the number and concentration of host strains (119, 120), and temperature (121–124). Furthermore, the addition of certain antibiotics or glycerol to the media can improve plaque formation and visibility by activating the SOS system in bacterial hosts (125). Some phages require specific cation conditions for adsorption, and the parameters used in this present study may not have been optimal for the selected pathogen hosts. Future studies should test the same samples using optimized methods with varying soft agar and ion concentrations, as divalent cations are crucial for phage DNA penetration into host cells (126, 127). These challenges highlight the need for method development and optimization when isolating phages from animal samples. Novel approaches should refine existing techniques by adjusting these critical factors to improve phage recovery. The lack of phages observed against specific pathogenic bacterial hosts in the present study highlights the need for further refinement of screening parameters, as the tested conditions may not be ideal to support optimal phage-host interactions.

## Conclusions

This study aimed to optimize the methodology for isolating bacteriophages from the bovine gastrointestinal, reproductive, and respiratory tracts, as well as cattle fetal fluid samples. While efforts were made to screen over 1,200 samples and isolate phages against bovine pathogens such as *T. pyogenes, M. haemolytica*, and *P. multocida*, the primary outcome of this research was the successful refinement of screening techniques, particularly Method 12. Despite encountering challenges in identifying phages against specific pathogens, we were able to isolate a significant number of phages targeting commensal bacteria, demonstrating the potential of optimized methods for phage discovery. The refined approach enhances the efficiency of phage isolation using traditional methods like the double-layer agar, which, despite being cost-effective and straightforward, requires adaptation for specific bacterial hosts and phage-host interactions. Through this optimized method, we successfully identified phages from diverse biological sources, including ruminal fluid, fecal, vaginal, nasopharyngeal, ocular, and fetal ruminal fluid. These phages exhibited lytic activity against 26 different bacterial genera that are commensal species originating from the rumen (majority), vagina, uterus, and intestinal tract of cattle. This highlights the potential of the cattle microbiome as a reservoir for phages with possible applications in microbiome modulation and disease control. Our study provides a basis in isolation and characterization of the bacteriophages from bovine gut, respiratory and reproductive tract, which is important in developing strategies to harness the power of bacteriophages for improved cattle health and feed efficiency.

## Author contributions

SA conceived the idea, secured funding, supervised the student, managed the project, assisted with sample collection, and critically revised the manuscript. GM designed and optimized the experimental methods, conducted sampling and data acquisition, and contributed to manuscript writing and editing.

## Institutional Review Board Statement

Not applicable.

## Informed Consent Statement

Not applicable.

## Data Availability Statement

All raw genome sequences and assemblies were deposited in the Sequence Read Archive (SRA) and GenBank, respectively, and can be accessed through the accession numbers are available from our two previous publications describing the genomes and genomic characteristics of the phages and their respective hosts (60, 76).

## Acknowledgments

The authors would like to thank Sarah Luecke, Emily Webb, Godson Aryee, Justine Kilama, Kell Hellmuth, and Kacie Smith in Amat lab for their assistance with sample collections. We also thank Dr. Barney Geddes and Tania Gupta at NDSU Microbiological Sciences for their contributions in perfecting the phage isolation protocols; Dr. T. G. Nagaraja from the KSU for kindly providing the *T. pyogenes* isolates; Dr. Kelli Maddock from the NDSU VDL for providing the *M. bovis* and *M. bovoculi* isolates; Dr. Heidi Pecoraro at the NDSU VDL and Marc Allard at NDSU Nebraska for providing liver abscess samples. Special thanks also to Scott Payne and Jayma Moore at the NDSU Electron Microscopy Core for their assistance with TEM imaging of the phages. We thank Scott Hoselton from NDSU Microbiological Sciences for his assistance in using the Inovaprep concentrating pipette, and for providing the wastewater samples; Jennifer Hulbert from NDSU Animal Sciences for her assistance with the biopsy and providing bovine liver tissue samples. We are grateful to Drs. Zack Carlson, Carl Dahlen, Joel Caton and Kendall Swanson from NDSU Animal Science for providing access to collect samples from their animal trials.

## Supplementary Materials

N/A

